# Exploration of the immune cell landscape in brain cancer utilizing gene expression and copy number data

**DOI:** 10.1101/490599

**Authors:** Yuriy Gusev, Krithika Bhuvaneshwar, Subha Madhavan

## Abstract

Brain cancer is a common cancer that affects more than 700,000 people in the US every year. We explore the dynamic changes in the abundance of immune cells based on RNA and DNA samples extracted from a large cohort of brain cancer patients. We used gene expression data and copy number data from a large brain cancer collections - the REMBRANDT project (REpository for Molecular BRAin Neoplasia DaTa) that includes 671 patients.

We applied virtual flow cytometry tools CIBERSORT and xCell to estimate the abundance of the immune cells in the RNA of these samples. The immune cell landscape in this dataset is compared with that of the TCGA brain cancer collection, that includes 511 patients with Lower Grade Glioma (TCGA-LGG) and 156 patients with Glioblastoma (TCGA-GBM).

We also discuss how well the results align with published literature, and how this computational analysis can help better understand how immune cells affect clinical outcome and survival in brain cancer patients

## INTRODUCTION

Brain cancer is a common cancer that affects more than 700,000 people in the US every year. Of these brain tumors, 80% are benign and 20% are malignant. More than any other cancer, brain cancers can have lasting life-altering impacts on a patient’s life. This is because brain cancers, including benign tumors, can interfere with portions of the brain responsible for viral body functions including speech, motor functions etc.

The average survival rate for malignant brain tumor patients is only 33.8% in men, and 36.4% in women. The survival rate for the most common malignant brain tumor - Glioblastoma (GBM) is only 5.5% ^1,2^.

A total of 130 sub types of brain cancers have been discovered. Some relevant to this dataset are described here. Most common primary brain tumor is called Meningioma. These rise in the meninges (brain lining), and mostly occur in the 70s or 80s. These are typically slow growing, and can be of tumor grade 1, 2, or 3. Astrocytoma are the cancer of the cerebrum, and can be of any tumor grade. High grade (grade 4) Astrocytoma is called Glioblastomas (GBM). Oligodendroglioma are cancer in the cells that make the covering that protects nerves. These are also typically slow growing, and can be of tumor grade 1, 2, or 3. For most brain tumor types, surgery and radiation remain the standard of care. There are only four approved drugs for brain tumors, and little has changed over the last 30 years^2^.

All these facts mentioned above are related to primary brain tumors. Secondary brain tumors are tumors that start elsewhere in the body and metastasize to the brain. Approximately 80% of cancers are known to metastasize in the brain^2^.

Brain tumors are known to have the highest cost of care for any cancer group^2^. This combined with limited treatment options makes brain cancers a deadly disease. It is hence imperative to improve the prognosis and treatment of this cancer group.

In recent years, immunotherapy has shown much promise in the treatment of various cancers. The microenvironment of the normal brain and early stage tumors is immunosuppressive due to the blood–brain barrier (BBB). The central nervous system (CNS) is known to be ‘immune privileged’ due to the BBB, which limits infiltration of molecules and helps regulation of immune cells normal circumstance^3^. However, this viewpoint of immune privilege has been revised. The antigens derived from CNS have been shown to induce immune response. In cancer, the BBB often gets compromised resulting in more infiltration of immune cells^3,4^.

In this analysis, we explore the dynamic changes in the expression of immune cells in the RNA of brain cancer patients. We used gene expression data from one of the largest brain cancer collections - the REMBRANDT project (REpository for Molecular BRAin Neoplasia DaTa) project that includes 671 patients^5,6^. We applied virtual flow cytometry tools CIBERSORT^7^ and xCell^8^ to estimate the abundance of the immune cells in the RNA of these samples. We examine the immune cell landscape in this dataset and compared with that of the TCGA brain cancer collection, which includes 511 patients with Lower Grade Glioma (TCGA-LGG) and 156 patients with Glioblastoma (TCGA-GBM). We also discuss how well the results align with published literature, and how this computational analysis can help better understand how immune cells affect clinical outcome and survival in brain cancer patients

## DATA

The dataset used in this analysis is called the **REMBRANDT dataset** (REpository for Molecular BRAin Neoplasia DaTa). It is a large brain cancer cohort, that includes 671 patients collected from 14 contributing institutions from 2004-2006^5,6^. The project won the Service to America Award in 2005^9^. Madhavan et al^6^ demonstrated the power of the data portal through several case studies.

The dataset includes a total of 671 patients with clinical data, of which 541 had gene expression data, and 507 patients had undergone SNP chip profiling. 263 patients had information about segment level copy number data. 220 patients had both gene expression and copy number data. Such combined datasets would provide researchers with a unique opportunity to conduct integrative analysis of gene expression and copy number changes in patients alongside clinical outcomes (overall survival).

For this analysis, we wanted to compare the immune infiltrates between the disease sub-groups. **Table 1** shows a summary of various disease groups available in this data set

**Table 1:** Summary of various disease groups in the REMBRANDT dataset.

| Clinical Attribute |  | Number of patients | % Of patients |
| --- | --- | --- | --- |
| Disease Type | GBM | 261 | 38.9 % |
|  | Astrocytoma | 170 | 25.3 % |
|  | Oligodendroglioma | 86 | 12.8 % |
|  | Non tumor | 31 | 4.6 % |
|  | Unknown | 68 | 10.1 % |
|  | Unclassified | 1 | 0.1 % |
|  | Mixed | 13 | 1.9 % |
|  | Blank/NA | 41 | 6.1 % |

## METHODS

We used the REMBRANDT dataset to explore the dynamic changes in the expression of immune cells. We first applied virtual flow cytometry tools CIBERSORT^7^ and xCell^8^ on the processed gene expression data to estimate the abundance of the immune cells in the RNA of these samples. The raw gene expression data in the form of ‘.CEL’ files, were initially processed according to the instructions of the CIBERSORT tool.

The estimated immune cells output was visualized using stacked bar graph and box plots. This allowed a visual comparison of the immune cell landscape of this dataset with that of the TCGA brain cancer collection and other public datasets. Gentles et al^7^, the creators of the CIBERSORT tool, applied their tool to the TCGA cancer collection that included brain cancers and other public datasets (https://precog.stanford.edu/iPRECOG.php)^7^.

We then used the estimated immune cells output to compare various disease sub-groups. We focused on the comparison between Astrocytoma (Astro), Glioblastoma and Oligodendroglioma (Oligo). We used non parametric Wilcoxin Test^10^ to compare each of these two groups. The results were in the form of differentially changed immune cell types.

We also compared these three groups in terms of copy number data, which is the form of chromosome instability index (CINdex). In Gusev et al^5^, we described the processing steps for this data conversion which was done using the CINdex Bioconductor package^11^. The CINdex package uses the segment level data to calculate the genomic instability in terms of copy number gains and losses separately at the chromosome and cytoband level. The genomic instability across a chromosome offers a global view (referred to as Chromosome CIN), and the genomic instability across cytobands regions provides higher resolution (referred to as Cytobands CIN) view of instability. This allows assessing the impacts of copy number alternations on various biological events or clinical outcomes by studying the association of CIN indices with those events. Each of these analyses in G-DOC was done on all patients that had data available.

This comparison of the disease sub-groups using the CINdex data was performed using the G-DOC platform using a student’s T test^12^. The results were in the form of differentially changed cytobands. We then used the CINdex Bioconductor package to find out which genes were present in these differentially changed cytoband regions. After we obtained the gene list, we performed system biology analysis to find out which biological processes were implicated (using R package EnrichR). **Figure 1** shows the workflow diagram

**Figure 1:**
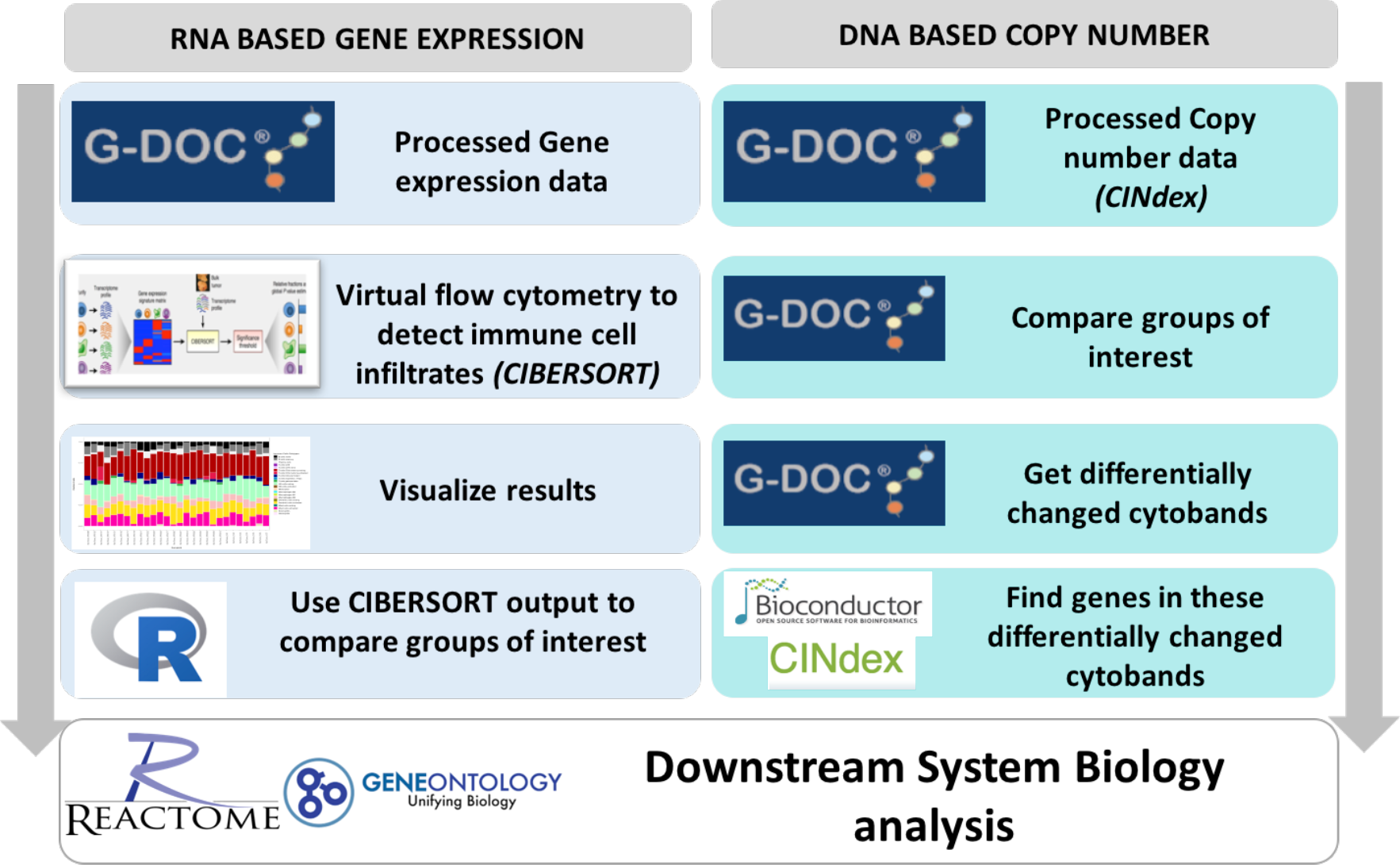
Analysis workflow.

## RESULTS

### Analysis of immune cell landscape based on gene expression data

**Figure 2A** shows the immune cell landscape of the Rembrandt data using estimation from CIBERSORT. **Figure 2B** is from Gentles et al (2015)^7^ showing immune cell landscape of the TCGA data collection (also using CIBERSORT). **Figure 2C** shows immune cell landscape of various public datasets (also using CIBERSORT), taken from iPRECOG^7^.

**Figure 2:**
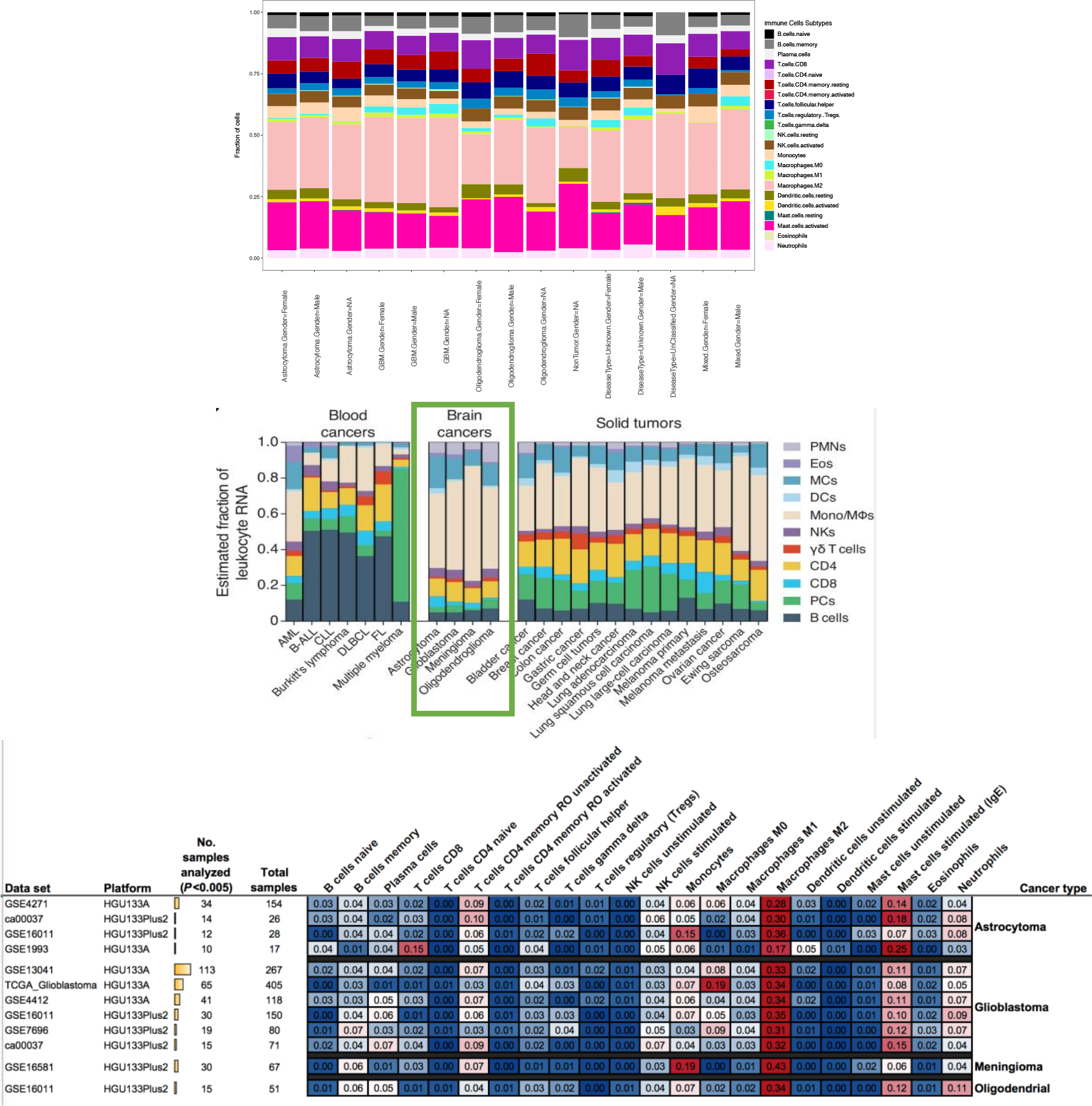
Figure 2A shows the immune cell landscape of the Rembrandt data using estimation from CIBERSORT. Figure 2B is from Gentles et al (2015) [ref] showing immune cell landscape of the TCGA data collection (also using CIBERSORT). Figure 2C shows immune cell landscape of various public datasets (also using CIBERSORT), taken from iPRECOG website.

We also visualized the estimated immune cell types using box plots. **Figure 3A** shows the box plot of immune cells estimates from the tool CIBERSORT, while **Figure 3B** shows immune cell fractions estimated by the tool xCell.

**Figures 3A and 3B:**
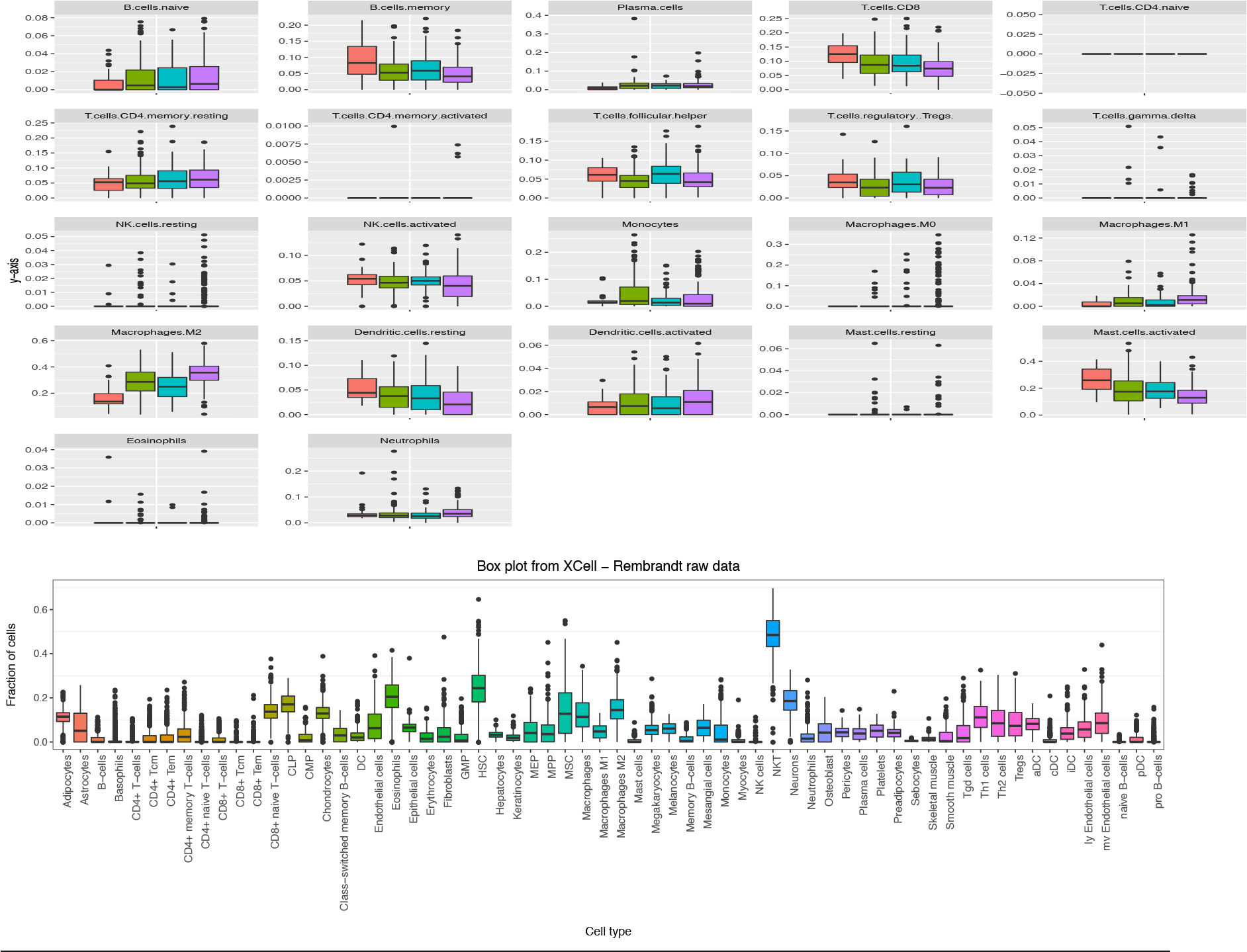
Box plot of immune cells from CIBERSORT and xCell respectively.

#### Comparison of immune infiltrates in several types of brain cancer

We also used the estimated immune cells output from CIBERSORT to compare various disease sub-groups - Astrocytoma (Astro), Glioblastoma and Oligodendroglioma (Oligo) using non-parametric Wilcoxin Test. The differentially changed immune cell type results are shown in **Table 2A, B** and **C**.

**Table 2A:** GBM vs. Astrocytoma.

| Feature | SignedFC | W statistic | P-value | Mean of CompGroup | Mean of BaseGroup | FDR |
| --- | --- | --- | --- | --- | --- | --- |
| Macrophages.M0 | Inf | 2516 | 0.001 | 0.035 | 0.000 | 0.01 |
| Macrophages.M2 | 1.20 | 2736 | 0.001 | 0.363 | 0.302 | 0.01 |
| Macrophages.M1 | 1.58 | 2641 | 0.005 | 0.016 | 0.010 | 0.03 |
| Monocytes | -1.70 | 1397 | 0.005 | 0.034 | 0.057 | 0.03 |
| Dendritic.cells.resting | -1.48 | 1488.5 | 0.017 | 0.024 | 0.036 | 0.07 |
| Mast.cells.activated | -1.37 | 1522 | 0.026 | 0.131 | 0.179 | 0.08 |
| T.cells.CD8 | -1.20 | 1525 | 0.027 | 0.075 | 0.091 | 0.08 |

**Table 2B:** GBM vs Oligodendroglioma.

| Feature | SignedFC | W statistic | P-value | Mean of CompGroup | Mean of BaseGroup | FDR |
| --- | --- | --- | --- | --- | --- | --- |
| Macrophages.M2 | 1.57 | 2313 | 4.17E-09 | 0.363 | 0.232 | 8.76E-08 |
| Mast.cells.activated | -1.58 | 682 | 2.53E-04 | 0.131 | 0.206 | 0.003 |
| Macrophages.M1 | 2.09 | 1796 | 0.004 | 0.016 | 0.008 | 0.025 |
| T.cells.follicular.helper | -1.40 | 824 | 0.005 | 0.050 | 0.070 | 0.025 |
| Neutrophils | 1.57 | 1760 | 0.008 | 0.042 | 0.027 | 0.031 |
| NK.cells.activated | -1.37 | 860 | 0.009 | 0.039 | 0.053 | 0.031 |
| T.cells.regulatory..Tregs. | -1.69 | 889 | 0.014 | 0.026 | 0.044 | 0.043 |
| NK.cells.resting | 12.16 | 1607 | 0.017 | 0.005 | 0.000 | 0.045 |

**Table 2C:** Astrocytoma vs Oligodendroglioma.

| Feature | SignedFC | W statistic | P-value | Mean of CompGroup | Mean of BaseGroup | FDR |
| --- | --- | --- | --- | --- | --- | --- |
| T.cells.follicular.helper | 1.59 | 655 | 0.002 | 0.070 | 0.044 | 0.03 |
| Macrophages.M2 | -1.30 | 256 | 0.005 | 0.232 | 0.302 | 0.05 |

### Analysis of immune response landscape based on copy number data

#### Comparing groups using cytoband level CIN data

We also compared these three groups in terms of copy number data, which is the form of chromosome instability index (CINdex) using Students T-test using G-DOC. The differentially changed cytobands have been summarized in **Table 3** and visualized as a circus plot (Figure 4). Complete results from the T-test are available as **Supplementary File 1**.

**Table 3:** Comparing group using copy number data (CINdex cytoband data)

| Comparison | # of differentially changed cytobands | # genes in these cytoband regions |
| --- | --- | --- |
| GBM vs Astrocytoma | 67 cytobands covering: 7p, 7q, 8q, 10q, etc | 1309 genes |
| GBM vs. Oligodendroglioma | 166 cytobands covering: 1p, 4q, 7p, 7q, 10q, 11p, 11q, 13q, 19q, etc | 4280 genes |
| Astrocytoma. Vs. Oligodendroglioma | 74 cytobands covering: 1p, 19q, etc | 2465 genes |

**Figure 4:**
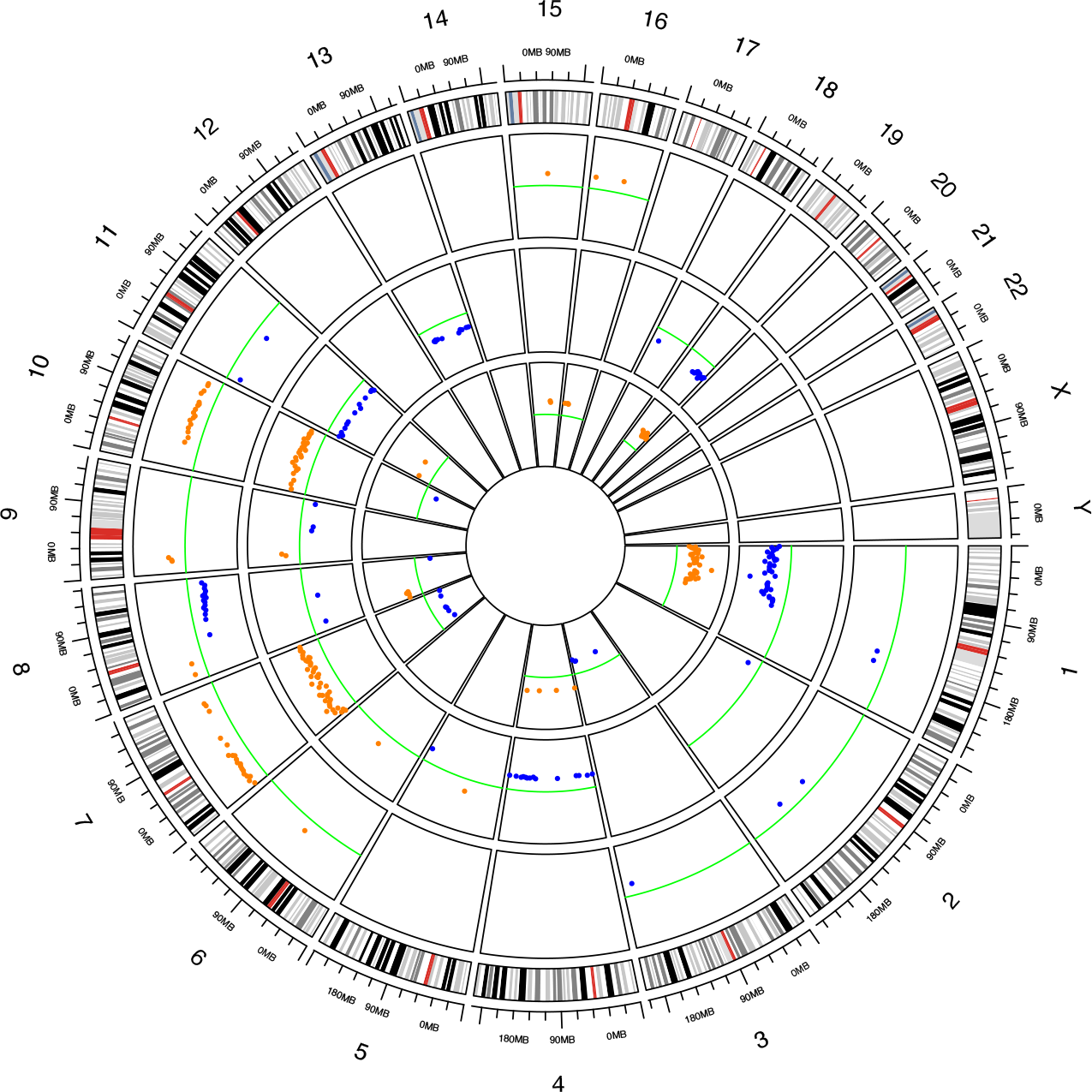
Circos plot showing significantly changed cytobands in each comparison. Each dot is a cytoband. The outer most track shows the significantly changed cytoband regions in the GBM vs. Astro copy number CIN data comparison. The middle track shows the significantly changed cytobands in the GBM vs. Olilgo CIN data comparison. The inner most track shows the significantly changed cytobands for the Oligo vs Astro CIN data comparison. The green line indicates 0 fold change. Cytobands with fold change > 0 are in orange dots, and those < 0 are in blue dots.

Once we obtained the genes located in the differentially changed cytoband regions, we performed system biology analysis. The biological processes enriched in Gene Onotology are shown in **Table 4**.

**Table 4:** Gene Ontology – biological processes enriched with genes located on cytobands with significantly different level of CIN index.

| GBM VS ASTROCYTOMA | GBM VS OLIGODENDROGLIOMA | ASTROCYTOMA. VS. OLIGODENDROGLIOMA |
| --- | --- | --- |
| signal transduction (GO:0007165) | regulation of transcription, DNA-templated (GO:0006355) | regulation of transcription, DNA-templated (GO:0006355) |
| natural killer cell activation involved in immune response (GO:0002323) | positive regulation of peptidyl-serine phosphorylation of STAT protein (GO:0033141) | phosphatidylinositol acyl-chain remodeling (GO:0036149) |
| positive regulation of peptidyl-serine phosphorylation of STAT protein (GO:0033141) | natural killer cell activation involved in immune response (GO:0002323) | neural tube closure (GO:0001843) |
| T cell activation involved in immune response (GO:0002286) | B cell differentiation (GO:0030183) | xenobiotic catabolic process (GO:0042178) |
| B cell differentiation (GO:0030183) | T cell activation involved in immune response (GO:0002286) | positive regulation of protein targeting to mitochondrion (GO:1903955) |
| response to exogenous dsRNA (GO:0043330) | response to exogenous dsRNA (GO:0043330) | liver development (GO:0001889) |
| humoral immune response (GO:0006959) | blood coagulation (GO:0007596) | epoxygenase P450 pathway (GO:0019373) |
| B cell proliferation (GO:0042100) | signal transduction (GO:0007165) | nuclear-transcribed mRNA catabolic process, nonsense-mediated decay (GO:0000184) |
| regulation of type I interferon-mediated signaling pathway (GO:0060338) | B cell proliferation (GO:0042100) | multicellular organism development (GO:0007275) |
| blood coagulation (GO:0007596) | humoral immune response (GO:0006959) | negative regulation of autophagy (GO:0010507) |
| adaptive immune response (GO:0002250) | epoxygenase P450 pathway (GO:0019373) | negative regulation of gene expression (GO:0010629) |
| innate immune response (GO:0045087) | type I interferon signaling pathway (GO:0060337) | female pregnancy (GO:0007565) |
| type I interferon signaling pathway (GO:0060337) | retinoid metabolic process (GO:0001523) | apoptotic process (GO:0006915) |
| sensory perception of smell (GO:0007608) | regulation of type I interferon-mediated signaling pathway (GO:0060338) | miRNA mediated inhibition of translation (GO:0035278) |
| cytokine-mediated signaling pathway (GO:0019221) | steroid metabolic process (GO:0008202) | positive regulation of p38MAPK cascade (GO:1900745) |
| positive regulation of cell proliferation (GO:0008284) | cell cycle arrest (GO:0007050) | pre-miRNA processing (GO:0031054) |
| social behavior (GO:0035176) | response to virus (GO:0009615) | mismatch repair (GO:0006298) |
| negative regulation of transcription, DNA-templated (GO:0045892) | adaptive immune response (GO:0002250) | cell surface receptor signaling pathway (GO:0007166) |
| positive regulation of insulin secretion (GO:0032024) | xenobiotic catabolic process (GO:0042178) | regulation of lamellipodium assembly (GO:0010591) |
| glucose homeostasis (GO:0042593) | mismatch repair (GO:0006298) | interstrand cross-link repair (GO:0036297) |
| response to virus (GO:0009615) | positive regulation of transcription regulatory region DNA binding (GO:2000679) |  |
|  | negative regulation of transcription, DNA-templated (GO:0045892) |  |

## DISCUSSION

From **Figure 2**, we can see that majority of the immune cells within brain tumors are macrophages. In **Figure 3A**, we can see that some cell types show immune suppression in higher grade cancer subtype compared to lower grade sub-types (e.g. B cells memory, T cells CD8, Dendritic cells resting, Mast cells activated). In the same figure, we can see that other cells types are upregulated in higher grade disease sub-types (E.g. Macrophages M2, CD4 memory resting, B cells memory naïve).

From the circus plot (**Figure 4**), we can easily spot the 1p/19q co-deletion in the group comparisons where Oligodendroglioma patient are involved. The 1/19q co-deletion has been used as a prognostic and predictive biomarker in Oligodendrogliomas ^13–15^.

In our results, Natural Killer (NK) cells were activated in GBM when compared to Oligo in both CIBERSORT and Copy number results. This is consistent with literature. It is known that the function of NK cells is often affected in brain cancer patients. One such example is TGFB down-regulates the expression of NKG2D activating receptor on NK cells in GBM patients. Decreased numbers of NK cells were observed in the blood of GBM patients post radiation treatment^16^.

In our data, we saw that GBM patients had high T cells CD4 memory resting, and lowest CD8 T cells (Figure 3A). According to published literature, GBM patients a high level of CD4+ TILs combined with low CD8+ TILs was associated with unfavorable prognosis ^17^.

Tumor aneuploidy (somatic copy number alterations) correlates with markers of immune evasion and with reduced response to immunotherapy^18^. Highly aneuploid tumors show reduced expression of CD8+ T cells. In our results, we see reduced T cells CD8 in GBM (**Figure 3**). Hence results are consistent with literature.

Also, tumor associated macrophages (TAMs) within the brain tend to be pro-tumorigenic and accumulate with higher tumor grade. TAMs have been implicated in brain tumor angiogenesis and resistance to anti-angiogenic therapies^4^.

It is important to note that copy number results from our analysis suggest similar pattern of immune cell response as Cibersort results. We found that the majority of categories of biological processes that are affected by cytoband level instability are mostly related to the processes involving the same categories of immune cells such asL B cells proliferation and differentiation, T cells activation, Natural Killer Cells activation etc.. These findings indicate that copy number changes in brain tumors playing important role in affecting immune cell activity in brain tumor microenvironment.

### Tumor microenvironment in brain cancers

The microenvironment of the normal brain & early-stage brain tumors is generally immunosuppressive due to the blood–brain barrier (BBB). Immune privilege qualities of the Central Nervous System (CNS) have been attributed to the BBB which limits transit of molecules and helps regulate lymphocyte tracking under normal circumstance^3^. However, this viewpoint of immune privilege has been revised. The CNS derived antigens have been shown to induce immune response. In cancer, the BBB often gets compromised resulting in leakiness^3,4^.

### Recent therapies

Some recent therapies immuno-therapies in brain cancer are listed below. These include (a) To develop strategies that re-educate macrophages to specifically adopt anti-tumor phenotypes in cancer. (b) Enhancing T cell activation by enabling co-stimulation, e.g., through the use of checkpoint inhibitors (c) Dendritic cells vaccines are likewise gaining significant clinical attention as an alternative strategy to stimulate T cell responses and (d) Monocyte/macrophage activation test has been developed as a cellular test of diagnostics and therapy^19^.

## DATA AVAILABILITY

The Rembrandt dataset is accessible for conducting clinical translational research using the open access Georgetown Database of Cancer (G-DOC)^20,21^ platform (https://gdoc.georgetown.edu). In addition, the raw and processed genomics and transcriptomics data have also been made available via the public NCBI GEO repository as a super series GSE108476. The MRI medical images for this dataset is available via The Cancer Imaging Archive (TCIA) initiative (https://wiki.cancerimagingarchive.net/display/Public/REMBRANDT).

